# Infection-specific long-chain fatty acid metabolism as a broad anti-enterovirus target

**DOI:** 10.64898/2026.06.10.731372

**Authors:** Marcos Florentin, Jeffrey Loube, Ekaterina G. Viktorova, Samuel Gabaglio, Elizabeth Tanner, Margaret A. Scull, George A. Belov

**Author notes:** Corresponding author: 8075 Greenmead Dr., College Park, MD, 20742.

## Abstract

Enteroviruses are arguably the most numerous group of viruses infecting humans. While most enterovirus infections are benign and self-resolving, their sheer number inevitably increases the chances of multiple complications. The diversity of enteroviruses means that the development of vaccines is only economically feasible against a select few, and no direct-acting or host-targeted anti-virals are approved to treat enteroviral infections, largely due to the rapid development of resistance against all experimental drugs. Here, we explored a universal property of enterovirus infection – a massive upregulation of phospholipid synthesis as a target for anti-viral interventions. The increased phospholipid synthesis consumes endogenously- and exogenously-derived long-chain fatty acids (LCFA). We demonstrate that polyunsaturated LCFAs can have a broad anti-enteroviral effect, affecting multiple steps of the virus life cycle. The anti-viral activity of LCFAs did not strictly depend on the degree of unsaturation or their capacity to induce lipid peroxidation but significantly correlated with their conformation. This suggests that their incorporation into the phospholipid molecules makes the replication organelle membranes incapable of properly accommodating viral replication machinery. Accordingly, the inhibition of neutral lipid synthesis promoted LCFAs retargeting to the membranes in infected cells and increased their anti-viral potency. We show that this approach is effective against diverse enteroviruses in different cell types, including differentiated primary cells, and that attempts to establish viruses resistant to such treatment were unsuccessful.

## Introduction

Enteroviruses are small non-enveloped viruses with a single-stranded positive RNA genome. A recently updated taxonomy recognizes 15 enterovirus species within the *Enterovirus* genus of the *Picornaviridae* family. Among them, *E. alphacoxsackie, betacoxsackie, coxsackiepol, deconjuncti*, *alpharhino*, *betarhino*, and *cerhino* species contain more than 300 distinct genotypes of human-specific viruses. While most enterovirus infections are benign and self-resolving, they can induce significant short- and long-term complications, such as partial or full paralysis, meningitis, aseptic encephalitis, acute and chronic myocarditis, and could trigger the development of type I diabetes (1–3). Only poliovirus and enterovirus A71 (members of the *E. coxsackiepol* and *alphacosackie* species, respectively) can be currently controlled with vaccines, and no specific anti-enterovirus drugs are approved for clinical practice despite a long history of developing compounds that target either viral proteins or host factors hijacked for the viral life cycle (reviewed in (4, 5)). The experimental drugs that reached the clinical trial stage were not advanced further because of the development of resistant variants and/or lack of clinical efficacy (6–9).

A hallmark of enterovirus infection is the profound reorganization of the intracellular membrane architecture, resulting in the development of complex, specialized membranous structures called replication organelles. The replication organelles provide optimal lipid and protein composition for the assembly and functioning of the viral replication complexes, and also shield dsRNA, an inevitable intermediate of RNA virus replication, from cellular sensors (reviewed in (10–12)). Replication organelles may occupy virtually the entire cellular volume by the end of the enterovirus replication cycle, and their structural development relies on the rapid activation of phospholipid, in particular, phosphatidylcholine, synthesis (13–16). Upregulation of phospholipid synthesis requires an increased supply of long-chain fatty acids (LCFAs) that constitute the hydrophobic part of phospholipid molecules. In principle, cells have access to three sources of LCFAs that can sustain new phospholipid synthesis: neutral lipid hydrolysis, such as triglycerides and cholesteryl esters stored in lipid droplets (LDs), *de novo* synthesis by fatty acid synthase (FASN), or import from the extracellular medium. In non-infected cells, the fine-tuned regulation of production, re-utilization, and degradation of LCFAs maintains the dynamic equilibrium of the cellular organelle membranes and allows metabolic adaptation in response to extracellular and intracellular cues. The relative contribution of different LCFA sources likely varies in different cell types, especially *in situ* in different tissues, depending on the extracellular milieu, the metabolic state, and the gene expression profile of given cells. In the case of poliovirus infection, at least in conditions of cell culture, it has been shown that LCFAs released from LDs support the bulk of infection-specific activation of phospholipid synthesis (17, 18).

The composition of LCFA moieties in the phospholipid molecules defines the basic physico-chemical properties of the membrane bilayer, such as its thickness and fluidity, which affect the recruitment and structural conformation of membrane-interacting proteins, contributing to the regulation of their functioning. In addition, polyunsaturated LCFA moieties in the membrane bilayer can undergo iron-dependent peroxidation at the bis-allylic position (19). The accumulation of lipid peroxides beyond a certain threshold leads to their uncontrollable propagation, triggering irreversible membrane damage and cell death. Such iron-dependent cell death, termed ferroptosis, is an important component of organismal homeostasis, including anti-pathogen responses, and can be actively manipulated by viruses (reviewed in (20)).

To enter metabolic pathways, LCFAs need to be thioesterified to coenzyme A (CoA) by long-chain acyl-CoA synthetase enzymes (ACSL). While LCFAs are hydrophobic molecules and can diffuse through the cellular membrane in both directions, acyl-CoAs are hydrophilic and cannot exit the cytoplasm (21, 22). Thus, the activation of ACSLCs upon infection, required to support the increase of phospholipid synthesis, also results in the activation of LCFA import from the extracellular medium (16).

In this study, we explored the possibility of manipulating the LCFA composition of replication organelle membranes as a universal anti-enterovirus strategy. We investigated the effect of LCFAs with a C18 backbone but with different degrees of unsaturation on the enterovirus replication cycle, and found that their anti-viral effect does not correlate with the capacity to induce lipid peroxidation, but rather with their conformation, suggesting that specific physicochemical properties of the phospholipid bilayer are important for assembly and/or functioning of the viral replication complexes. We identified acyl-CoA cholesterol transferase activity (ACAT) as the primary activity responsible for upregulation of neutral lipid synthesis upon short-term exposure of cells to the increased concentration of LCFAs. Accordingly, inhibition of this activity strongly increases the anti-viral potency of LCFAs, consistent with promoting their routing into the phospholipid synthesis. Importantly, we show that LCFAs can similarly inhibit the replication of diverse enteroviruses in different cell types, including a physiologically relevant infection model, and that selection of resistant mutants to such inhibition was not successful.

Thus, our data provide important insights into the virus-cell interaction and demonstrate the potential of LCFA-like molecules as a novel class of broad anti-viral compounds with a high barrier to the development of resistance.

## Materials and Methods

### Cells and viruses

Human cervical carcinoma HeLa cells were a gift from Dr. Ehrenfeld, NIH; human lung carcinoma A549 and human rhabdomyosarcoma RD cells were purchased from ATCC. HeLa and RD cells were grown in DMEM high-glucose modification supplemented with pyruvate; A549 cells were grown in F12K medium. All growth media were supplemented with 10% heat-inactivated FBS, 1x penicillin-streptomycin solution, and 5 mg/mL Plasmocin. Poliovirus (PV) type 1 strain Mahoney was a gift from Dr. Ehrenfeld, NIH; Coxsackievirus B3 (CVB3) strain Nancy was rescued from the plasmid p53CB3T7, kindly provided by Professor Frank van Kuppeveld (Utrecht University, the Netherlands); Rhinovirus A16 (RVA16) (23) was rescued from a cDNA clone kindly provided by Dr. Ann Palmenberg (University of Wisconsin-Madison). Enterovirus D68 (EVD68) strain USA/2018-23088 was generously provided by Dr. Amy Rosenfeld, FDA. Infections with all enteroviruses in cell culture were performed similarly. Upon removal of growth medium, cell monolayers were washed once with fresh serum-free growth medium, and upon the addition of the viral inoculum, the cells were incubated with soft rocking at room temperature for 30 min (PV and CVB3) or 60 min (EVD68). The viral inoculum was removed, and the medium with the indicated compounds was added. Infected cells were incubated at either 37 °C (PV, CVB3) or 33 °C (EVD68) in 5% CO_2_. The total virus yield was determined upon freeze-thawing the infected cells as TCID_50_/ml using titrations on HeLa or RD cells grown in 96-well plates; the titers were calculated using Karber’s formula (24). Viral titers were determined from at least three biological replicates.

Replication of RVA16 in HAE cultures was assessed by qPCR. Briefly, total RNA was isolated using the RNEasy kit (#74106, Qiagen) following the manufacturer’s instructions, and eluted in 33 µL of water. cDNA was synthesized from 100 ng of total RNA with the High-Capacity cDNA Reverse Transcriptase kit (#4368814, Applied Biosystems) using random hexamer primers. Three microliters of cDNA were then combined with 10 pM of forward and reverse RV primers (5′–ACMGTGTYCTAGCCTGCGTGGC–3′, and 5′–GAAACACGGACACCCAAAGTGT–3′), 10 pM HRVTF probe (5′–FAM/TCCTCCGGCCCCTGAAT/BHQ1–3′), all acquired from Integrated DNA Technologies, and 7.5 µl LuminoCT Taqman Master Mix (#L6669, Sigma-Aldrich), in a final volume of 15 µl. Samples were run on a LightCycler 480 (Roche) in the Imaging Core Facility in the Department of Cell Biology and Molecular Genetics at the University of Maryland, College Park. Serial dilutions of RVA16 cDNA with known concentrations were run alongside experimental samples, and exponential regression on a semi-log scale of this standard curve was used to convert the cycle threshold (Ct) values to genome copy number for each sample.

### Human airway epithelial (HAE) cultures

Human tracheobronchial epithelial cells, sourced from three different donors without lung disease, were purchased from Lonza (#CC-2540S). HAE cells were expanded and then differentiated in parallel at air-liquid interface to yield a pseudostratified epithelium as previously described (25). Briefly, 6.5 mm Transwell membranes with 0.4 µm pores (# 3470, Corning) were coated with rat tail collagen type 1 (#354236, Corning), then 3.3E4 cells were seeded per Transwell in Pneumacult Ex-plus medium (#05040, Stemcell Technologies). Four days after seeding, the apical medium was aspirated, and the basolateral medium was replaced with PneumaCult-ALI medium (#05001, Stemcell Technologies). Cultures were maintained at 37 °C with 5% CO_2_, and allowed to differentiate for 4 weeks with basolateral medium changed every 2-3 days.

For infection with RVA16, the apical surface of HAE cultures was washed with 100 µl of pre-warmed PBS to remove mucus. 500 µl of PneumaCult-ALI and 10 µl of PBS with 200µM of LCFAs and 50µM of Avasimibe (ACAT inhibitor), or with the corresponding amount of DMSO for control samples, were delivered to the basolateral and apical compartments, respectively. 10µL of PBS containing 5E5 PFU of RVA16 was then added to the apical compartment, and cultures were subsequently maintained at 34 °C with 5% CO2. At ∼24 h p.i, 100 µL of PBS was added to the apical surface, and the cultures were incubated for 30 min at 34 °C. The PBS was then aspirated, and culture membranes were removed from the plastic Transwell supports and lysed in 350 µl RLT buffer (#79216, Qiagen) for downstream RNA isolation. Basolateral medium was collected for cytotoxicity assay.

### Antibodies and chemicals

Mouse monoclonal anti-PV 2C antibody described in (26) was a gift from Professor Kurt Bienz, University of Basel, Switzerland; rabbit polyclonal anti-PV 3D antibodies were described in (27). Anti-β-Actin−HRP antibody conjugate was from Sigma-Aldrich (#A3854). Stearic Acid (#10011298), Oleic Acid (#90260), Linoleic Acid (#90150), α-Linolenic Acid (#90210), γ-Linolenic Acid (#90220), α-Eleostearic Acid (#10008349), β-Eleostearic Acid (#22976), Punicic Acid (#26057) and 3(Z),6(Z),9(Z),12(Z),15(Z)-Octadecapentaenoic Acid (ODPA) (#10009731), were from Cayman Chemicals. ACAT1 and 2 inhibitor Avasimibe (#HY-13215), DGAT1 inhibitor T863 (#HY-32219), and DGAT2 inhibitor PF-06424439 methanesulfonate (#HY-108341A) were from MedChemExpress. BODIPY 500/510 C4, C9 (#B3824) and BODIPY 581/591 C11 (#D3861) were from Molecular Probes (Thermo Fisher). Stocks of fatty acids and inhibitors were prepared in DMSO to a final concentration of either 10 or 25 mM.

### Toxicity assays

Cell viability was measured using either the CellTiter-Glo Luminescent Cell Viability Assay (#G7570, Promega), which is based on the cellular ATP content, or the CytoTox 96 Non-Radioactive Cytotoxicity Assay kit (#G1780, Promega), which quantifies LDH release resulting from plasma membrane damage. For CellTiter-Glo assays, HeLa cells were seeded in 96-well plates at 3.5E4 cells/well. The following day, the medium was replaced with that containing LCFAs and inhibitors and incubated for the specified time according to the experiment. After equilibration at room temperature, CellTiter-Glo substrate was added, and luminescence was read within 15 min using a SpectraMax iD3 (Molecular Devices) microplate reader. For fatty acid pre-loading experiments, cells were incubated overnight in a standard growth medium supplemented with LCFAs, and processed as described the following day.

LDH release assay was used to evaluate the viability of HAE cultures after infection with RVA16. Briefly, each basolateral medium sample was assessed in three technical replicates of 50 µl. Background samples contained 50 µl of unconditioned PneumaCult-ALI medium, and total lysis control for each donor was measured from 50µl samples of the lysate obtained by adding 100 µl of 1x lysis buffer from the kit into the apical compartment of the HAE culture, followed by incubation for 45 min at 37 °C with 5% CO_2_. All control and experimental samples were processed in the same 96-well plate. 50µl of the CytoTox 96 reagent was added to each sample well for 30 mins at room temperature in the dark before adding 50 µl of stop solution from the kit. The signal was measured at 490 nm using a Multiskan Ascent (Labsystems) plate reader. The average background value was subtracted from each experimental well, and % cytotoxicity was calculated according to the manufacturer’s instructions.

Outer membrane damage was quantified using Sytox Green Nucleic Acid Stain (#S7020) from ThermoFisher. HeLa cells were seeded in 12-well plates at 4.5E5 cells/well. The following day, cells were washed once with fresh DMEM and infected with poliovirus at an MOI of 50. Cells were incubated in DMEM without FBS until 2 h p.i., and LCFAs were added directly to the medium to a final concentration of 200 uM. At 3.5 h p.i., Sytox Green and Hoechst 33342 (a cell-permeable nuclear stain) were added to a final concentration of 30 nM and 0.5 µg/mL, respectively. Live cells were imaged using a Zeiss Axiovert 200M fluorescence microscope operated by ZEN software (Zeiss). After imaging, cells were washed once with PBS and frozen at −80 °C until further processing for Western Blotting.

### Lipid peroxidation assay

Lipid peroxidation was measured using the BODIPY 581/591 C11 Lipid Peroxidation Sensor. Cells were seeded in 8-chamber µ-Slides (#80807, Ibidi) at 1.1E5 cells/well. The following day, medium was removed, cells were washed twice with DMEM without FBS, and incubated for 30 min in DMEM without FBS containing 1 µM BODIPY 581/591 C11. Cells were then washed with fresh DMEM and infected with PV at an MOI of 50, kept in DMEM without FBS for 2 h p.i., and then provided with new medium containing 200 µM LCFAs and incubated for two more hours. For the last 30 min of incubation, Hoechst 33342 was added to a final concentration of 0.5 µg/mL, and live cells were imaged with a Zeiss LSM 510 confocal microscope operated by ZEN software (Zeiss) with the emission wavelength of 591 nm and 509 nm for reduced (red) and oxidized (green) forms of BODIPY 581/591 C11, respectively.

### Lipid droplet staining

Cells grown on coverslips in 12-well plates were incubated in a standard growth medium supplemented with 200 µM of LCFAs, where indicated. The next day, cells were washed once with PBS and fixed with 4% PFA (#15710, Electron Microscopy Sciences) in PBS for 20 min, then washed twice with PBS for 5 min with soft rocking at room temperature. The PBS was replaced with an Oil Red O staining solution (0.5% of Oil Red O (#O0625, Sigma-Aldrich) in isopropanol diluted 3:2 in molecular grade water), and the samples were incubated for 15 min with soft rocking at room temperature protected from light. Hoechst 33342 was used as a nuclear stain. Cells were washed with 70% ethanol for 5 s, rinsed with molecular-grade water, and the coverslips were mounted with Fluoromount-G (#17984-25, Electron Microscopy Sciences). Cells were imaged using a Zeiss LSM 510 confocal microscope.

### Quantitative analysis of microscopy images

Analysis of microscopy images was performed with Cell Profiler 4.2.8 software (28) using custom pipelines built for the segmentation of nuclei, cytoplasm, and lipid droplets. All images from the same experiment were taken with the same microscope settings and processed with the same Cell Profiler pipeline. Analysis was performed on full-size images without rescaling.

### Modeling of fatty acid conformation

Molecular modeling was performed with RDKit version 2024.09.6 (29). Code and description are available at the Belov Lab repository (FA-geometry): https://github.com/belov-lab/FA-geometry. The exact version used in this study is archived on Zenodo, https://doi.org/10.5281/zenodo.18190122. The workflow uses RDKit 2024.09.6; ETKDGv3 for conformers; MMFF94 minimization and Python 3.x. See docs/FA-geometry-method-csv.md and docs/FA-geometry-method.md for details.

### Statistical analysis

Statistical analysis and graphical presentation of experimental data were performed with the GraphPad Prism-11 software package. Graphs show average values and standard deviation bars. An unpaired two-tailed t-test was used for pairwise comparison; comparison of multiple samples with a control was performed with a one-way ANOVA test with Dunnett correction for multiple comparisons, p ≤0.05 was considered significant. * indicate 0.01<p ≤ 0.05, ** indicate 0.001< p ≤ 0.01, *** indicate 0.0001<p ≤ 0.001, and **** indicate p ≤ 0.0001. Principal component analysis (PCA) of fatty acid conformational properties was performed with R (4.5.1) using RStudio (2026.1.1.403). Fatty acid three-dimensional properties and PCA results were plotted using R rgl and ggplot2 packages, respectively (30, 31).

## Results

### The anti-viral activity of polyunsaturated fatty acids correlates with their conformation

The capacity of picornavirus-infected cells to redirect LCFAs toward structural phospholipid synthesis prompted us to explore the possibility of selective targeting of infected cells by inducing lipotoxic stress. We hypothesized that increased incorporation of polyunsaturated fatty acids (PUFAs) into membrane phospholipids in infected cells may promote lipid peroxidation, eventually triggering ferroptotic cell death. We screened the effect of several LCFAs on cell viability and poliovirus replication using HeLa cells. We tested the following fatty acids, all containing a C18 backbone but with variations in the number and position of double bonds: stearic acid (C18:0), oleic acid (C18:1), linoleic acid (C18:2), α-linolenic acid (C18:3), γ-linolenic acid (C18:3), α-eleostearic acid (C18:3), β-eleostearic acid (C18:3), punicic acid (C18:3), and all-cis-3,6,9,12,15-octadecapentaenoic acid (ODPA) (C18:5) (Fig. 1A). All these fatty acids are derived from natural sources, further, linoleic and α-linolenic acids are essential fatty acids for humans. The poliovirus replication cycle in HeLa cells takes 6-8 h and is accompanied by high activation of structural phospholipid synthesis (13–16). To isolate the effect of the incorporation of exogenously added fatty acids into the membranes of the replication organelles from possible interferences with other steps of the viral life cycle, HeLa cells were infected with poliovirus at an MOI of 50 (to ensure synchronous infection) and incubated for 2 h in a serum-free medium before the addition of fatty acids. By 2 h p.i., virion penetration and uncoating is complete, and the development of replication organelles begins. At this stage, only occasional minor modifications of the intracellular membranes are detectable by electron microscopy (32–35). At 2 h p.i., the medium was changed to that containing increasing concentrations of LCFAs, up to 200 µM, which is a standard concentration of exogenous LCFAs used in many studies of lipid metabolism (36–38). The cells were incubated for an additional 2.5 h after the addition of fatty acids and then processed for analysis. Cell viability was assessed by Sytox Green staining, which detects plasma membrane damage (39), and the accumulation of the viral protein 2C served as a measure of viral replication.

**Figure 1.**
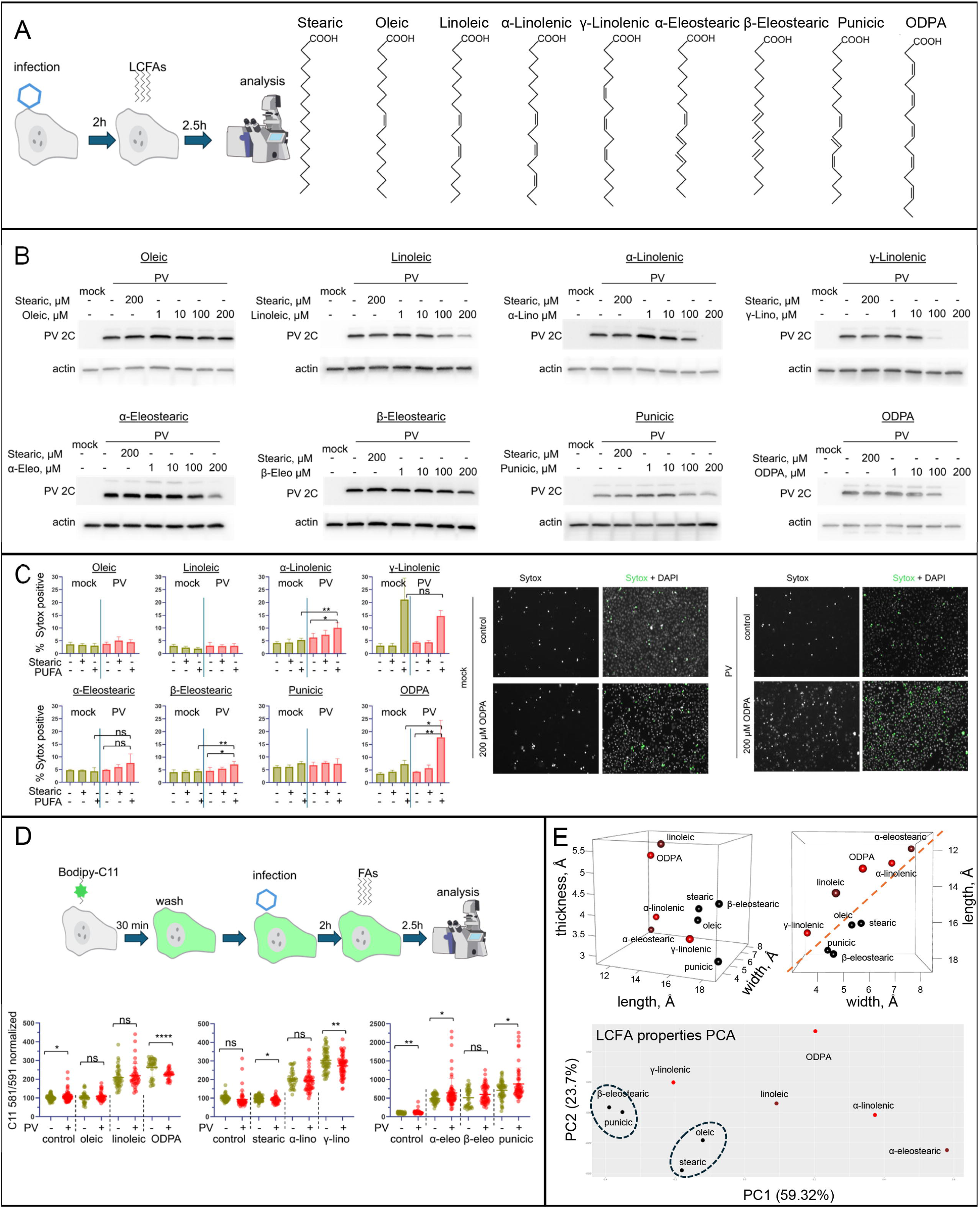
Anti-viral effect of polyunsaturated fatty acids correlates with their conformation. **A.** A scheme of the experiment for assessing the effect of exogenously added LCFAs in viral infection and bond-line structures of C18 LCFAs used in these experiments. **B.** Western blot analysis of the accumulation of poliovirus protein 2C in HeLa cells infected at an MOI of 50 and incubated in serum-free medium supplemented with the indicated LCFAs from 2 to 4.5 h p.i. Actin is shown as a loading control. **C.** Left panels: percentage of the Sytox green-positive nuclei indicating membrane damage in cells treated with 200 µM of the indicated LCFAs as in B. Quantifications for each sample were performed from four randomly chosen fields of view with an average of ∼1000 nuclei in each. Right panels: representative images of control cells and those incubated with ODPA showing Sytox green and DAPI signals used for quantifications. **D.** Top: a scheme of the experiment to measure the development of lipid peroxides using the C11 probe in cells treated with 200 µM of the indicated LCFAs after infection as in B. Bottom: quantitation of the C11 signal ratio at 581 nm (oxidized) to that of the original compound at 591 nm. Dots represent signals from individual cells. Data are normalized to the average signal in mock-infected control samples (i.e., incubated without addition of exogenous LCFAs). **E.** Top: two projections of the 3D-plotted length, width, and thickness of the predicted conformations of LCFAs used in these experiments. Black, brown, and red balls are assigned to LCFAs with no, moderate, and strong anti-viral activity as determined by the 2C accumulation in A, respectively. The red dashed line separates LCFAs without anti-viral activity on the bottom. Bottom: PCA analysis of the predicted conformations of LCFAs used in these experiments. Color coding is the same as in the top panels. Black ellipses outline LCFAs without anti-viral activity.

Saturated stearic acid and monounsaturated oleic acid did not show anti-viral activity even at the highest concentration tested; therefore, further on, stearic acid was used as a negative control for anti-viral effects in all following experiments. Surprisingly, we observed that the anti-viral activity of PUFAs did not strictly correlate with their degree of unsaturation, nor with the plasma membrane damage of infected cells. ODPA (C18:5), α-linolenic (C18:3), and γ-linolenic (C18:3) showed the strongest inhibition of viral replication, while the effects of α-eleostearic (C18:3) and linoleic (C18:2) were more modest, but also clearly detectable. At the same time, β-eleostearic (C18:3) and punicic (C18:4) acids did not significantly affect viral replication (Fig. 1B). The anti-viral effect of γ-linolenic acid was likely due to its high toxicity to both infected and non-infected cells, while ODPA, α-linolenic, and β-eleostearic were selectively more toxic to infected cells (Fig. 1C), yet the latter fatty acid did not affect viral replication.

To see if the incorporation of PUFAs into the membranes of replication organelles leads to increased lipid peroxidation, we monitored a fluorescence spectrum shift of a BODIPY C11 probe. The ratio of the fluorescence signal of the oxidized form of BODIPY C11 at 581 nm to that of the original compound at 591 nm provides a quantitative assessment of the accumulation of lipid peroxides, a standard technique in ferroptosis assays (40). BODIPY C11 is a long-chain fatty acid analog, so to ensure that infected cells and mock-infected controls incorporate the same amounts of the indicator molecule, the cells were pre-incubated with BODIPY C11 for 30 min, and the compound was removed before infection. The infection of BODIPY C11-prelabelled cells and incubation with 200 µM of LCFAs was performed similarly to the previous experiments (Fig. 1D). We observed a slight increase in lipid oxidation in control infected samples (i.e., incubated without additional LCFAs) in some but not all experiments. PUFAs similarly increased lipid peroxidation signals in both mock-infected controls and infected cells, although in some experiments, the signal could be statistically significantly higher in either case. For example, in the data presented in Fig. 1D, for ODPA and γ-linolenic acids, the signal was statistically significantly stronger in mock-infected cells, while in the case of α-eleostearic and punicic acids, the signal was stronger in infected cells (Fig. 1D). Thus, the anti-viral activity of C18 PUFAs does not correlate with their capacity to induce lipid peroxidation and cell death.

To gain insight into the mechanism of the anti-viral effect of LCFAs, we evaluated their conformations using RDKit modeling software (29, 41). This software generates several geometric measures of the ensemble of configurations the molecule is likely to assume under specific conditions. Interestingly, the anti-viral activity strongly correlated with the length and the width of LCFAs, so that those that did not affect viral replication were clearly separated from those that did, regardless of the degree of polyunsaturation. The analysis suggests that relatively wide and short LCFAs cannot be tolerated in the membranes of replication organelles (Fig. 1E, top). Only γ-linolenic acid does not seem to fall within this category, as it is relatively long and narrow, but this is likely due to the different mechanism of anti-viral activity related to γ-linolenic acid-induced toxicity in both infected and non-infected cells. The toxic effect of γ-linolenic acid on cancer cell lines has been observed previously (42, 43). The separation of LCFAs that did not affect viral replication also manifested in principal component analysis that takes into account additional conformational metrics of the LCFA molecules (Fig. 1E, bottom, Table S1).

Thus, exogenously added LCFAs can inhibit poliovirus replication, and their anti-viral activity is likely related to the modification of the properties of lipid bilayer upon their incorporation into phospholipid molecules, but not directly to the degree of unsaturation and induction of lipid peroxidation.

### Both exogenous and intracellular sources of long-chain fatty acids can affect viral replication

For further in-depth evaluations, we chose linoleic acid and ODPA, as a physiologically relevant essential LCFA, and a compound with a strong anti-viral activity, but which is not normally encountered by human cells, respectively.

We next set out to evaluate the anti-viral effects of exogenously-added PUFAs in the presence of serum. Incubation of HeLa cells in normal growth medium supplemented with up to 200 µM of stearic acid, linoleic acid, or ODPA for 6 h did not show any signs of toxicity. In contrast, the ATP-based cell viability assay demonstrated a statistically significant increase in signal for cells incubated with these LCFAs, likely reflecting their active metabolic targeting to oxidative phosphorylation and ATP production (Fig. 2A). To see if exogenous LCFAs in the presence of serum affect poliovirus yield during one replication cycle, HeLa cells were infected at an MOI of 5 or 0.5, and incubated in normal growth medium supplemented with 200 µM of LCFAs for 6 h. While a modest, within ∼0.5 log_10_, reduction of virus yield was observed in cells incubated with either linoleic acid or ODPA, this difference was not always statistically significant in experiments performed with three biological replicates (Fig. 2B).

**Figure 2.**
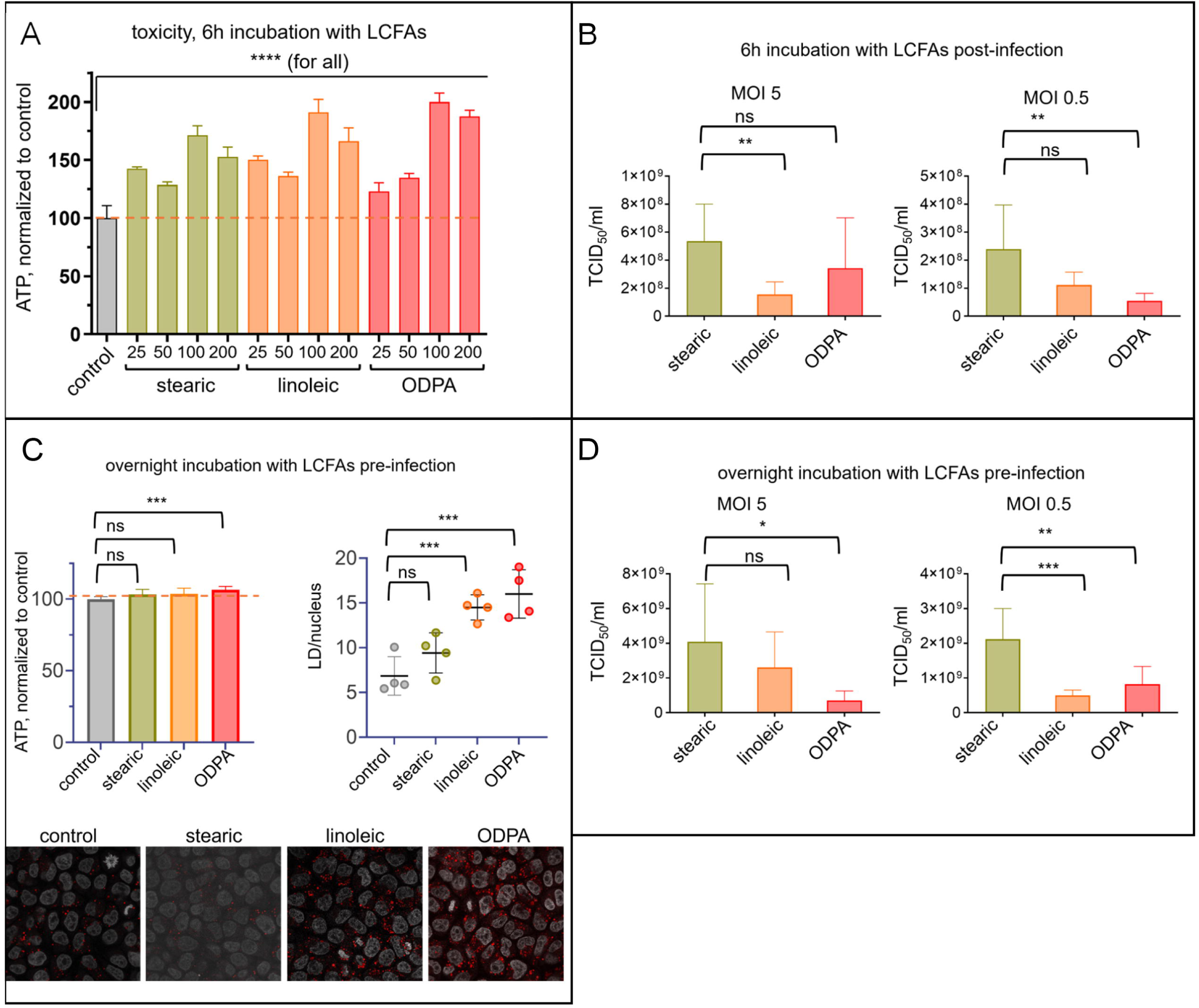
Effect of serum- and LD-derived LCFAs on poliovirus replication. **A.** Cytotoxicity assay (ATP level) of HeLa cells incubated in normal growth medium supplemented with the indicated LCFAs (concentration in µM is shown on X axis) for 6 h. **B.** Poliovirus yield in HeLa cells infected with an MOI of 5 or 0.5 and incubated after virus adsorption in normal growth medium supplemented with 200 µM of the indicated LCFAs for 6 h. Experiments were performed with three biological replicates. **C.** Top left: toxicity assay (ATP level) of HeLa cells incubated overnight in normal growth medium supplemented with 200 µM of the indicated LCFAs. Right: quantitation of LDs in these cells, each dot represents an individual randomly chosen field of view with ∼50 nuclei each. Bottom: representative images of Oil Red O staining of LDs used for quantitation. **D.** Poliovirus yield in HeLa cells incubated overnight in normal growth medium supplemented with 200 µM of the indicated LCFAs, infected with poliovirus at an MOI of 5 or 0.5, and incubated after the virus adsorption in a serum-free medium for 6 h p.i. Experiments were performed with three biological replicates.

Next, we assessed the effect of the composition of endogenously available fatty acids on virus replication. To preload cells with specific LCFAs, they were incubated overnight in normal growth medium supplemented with 200 µM of stearic acid (control), linoleic acid, or ODPA. Extended incubation of cells with 200 µM of LCFAs was also non-toxic, with the ATP signal showing minimal variation among samples (Fig. 2C, left panel). In cells incubated with linoleic acid and ODPA overnight, we observed a strong increase in the number of LDs, indicating that PUFAs are effectively targeted for neutral lipid synthesis and storage. In samples incubated with stearic acid, the increase in the number of LDs was less pronounced, but also noticeable, although it did not reach statistical significance (Fig. 2C, right and bottom panels). Upon infection, LCFAs stored in LDs would be mobilized and used for phospholipid synthesis, supporting the structural development of replication organelles (17, 18). Cells preincubated overnight with 200 µM of LCFAs were infected at an MOI of 5 or 0.5 of poliovirus and, to exclude the effect of exogenous fatty acids, incubated post-infection in a serum-free medium for 6 h (one replication cycle). As in the case with the exogenously added fatty acids, we observed a ∼0.5 to 1 log_10_ reduction of the virus yield in samples pre-loaded with linoleic acid and ODPA, even though the difference was not always statistically significant in experiments performed with three biological replicates (Fig. 2D).

Altogether, these data show that non-infected cells have effective short-term and long-term mechanisms to restore homeostasis in response to a significant increase in extracellular LCFA concentration, and that LCFAs imported from either the extracellular medium or mobilized from LDs can affect poliovirus replication.

### Neutral lipid synthesis in infected cells continues in the presence of PUFAs

Our results demonstrate that serum in the growth medium not only reduces the cytotoxic effect of PUFAs, but also significantly attenuates their anti-viral activity. Serum supplies a complex mixture of lipids that compete with exogenously added fatty acids and may activate alternative utilization pathways, thereby diverting them from incorporation into replication organelle membranes. This likely limits the impact of added PUFAs on the metabolism of infected cells.

To visualize the active fatty acid metabolism pathways, we incubated infected cells in a serum-supplemented growth medium containing 200 µM of stearic acid (control), linoleic acid, or ODPA for 4 h, and then added 0.4 µM BODIPY 500/510 C4, C9; a C18 fatty-acid analog with an internal BODIPY fluorophore, for 30 min (Fig. 3A). Hereafter, we will refer to this probe as a fluorescent fatty acid (FFA). After the incubation with FFA, live-cell fluorescence images were acquired. Thus, the intracellular distribution of the FFA reports on the LCFAs utilization pathways that are active during the 4.0-4.5h interval, in the middle of the replication cycle. This probe is broadly used for visualization of LCFA metabolism, and we have previously used this approach to analyze lipid synthesis during infection (16, 44, 45).

**Figure 3.**
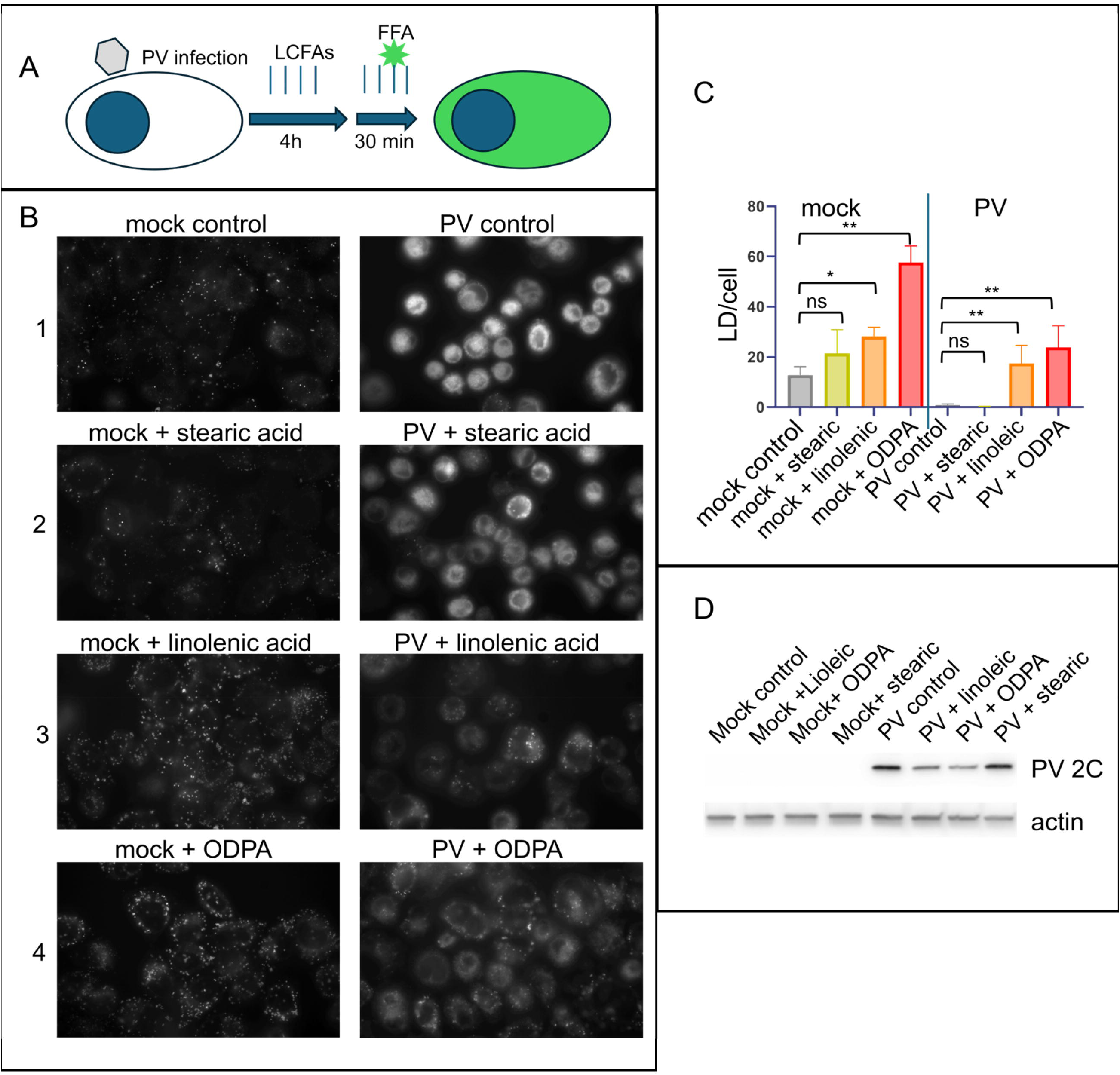
Lipid synthesis pathways in infected cells are affected by exogenously added LCFAs. **A.** A scheme of experiments to trace the active LCFA targeting pathways with the FFA probe. **B.** Representative images showing live HeLa cells infected with poliovirus with an MOI of 50 (or mock-infected controls)and incubated in the presence of 200 µM of the indicated LCFAs for 4 h plus 30 min with the same fatty acids and 0.4 µM of the FFA probe. The fluorescent signal shows cellular structures where FFA is incorporated during 30 min of incubation. **C.** Quantitation of LDs in cells treated as in A. At least ∼40 cells are quantified for each sample. **D.** Western blot with lysates of cells infected and incubated with LCFAs as in B. Actin is shown as a loading control.

In mock-infected cells incubated without exogenous LCFAs, the FFA signal localized predominantly to LDs, with only a weak staining of cytoplasmic membranous structures, while in infected cells incubated without added LCFA, the fluorescent signal was markedly stronger, and the FFA was actively incorporated into replication organelle membranes. The increase in FFA signal during infection reflects a strong upregulation of LCFA import and membrane phospholipid synthesis, consistent with prior observations (16) (Fig. 3B, panel 1). Mock and infected cells treated with stearic acid displayed FFA distributions closely resembling respective controls (Fig. 3B, panel 2). In contrast, in mock-infected cells incubated with PUFAs, the number of FFA-positive LDs per cell increased substantially, from ∼10 in controls to ∼20 with linoleic acid and ∼60 with ODPA, indicating robust activation of neutral-lipid synthesis (Fig. 3B, panels 3 and 4, mock; Fig. 3C). Notably, in infected cells incubated with linoleic acid or ODPA, FFA signal in replication organelle membranes was weaker than in infected controls, and targeting of FFA to LDs remained evident, whereas LD signal was virtually absent in infected controls (Fig. 3B, compare infected cells from panels 3 and 4, with those in panels 1 and 2; Fig. 3C). Western blot analysis of the viral protein 2C confirmed a moderate inhibition of viral replication by both PUFAs in these conditions, consistent with previous experiments shown in Fig. 3B.

These observations suggest that the strong activation of neutral lipid synthesis diverts PUFAs from phospholipid synthesis, attenuating their effect on viral replication. Therefore, we hypothesized that blocking the synthesis of neutral lipids may increase LCFA targeting to the membranes of replication organelles and improve their anti-viral efficacy.

### Acyl-CoA cholesterol acyltransferase is the major activity driving the activation of lipid droplet formation upon exposure to polyunsaturated fatty acids

LD cores mainly consist of triglycerides and cholesteryl-LCFA esters (reviewed in (46, 47)), therefore, the FFA signal in LDs may reflect the activation of synthesis of either of these neutral lipids. The final step in the major pathway of triglyceride synthesis is the addition of a long-chain acyl moiety to the 1,2-diacylglycerol backbone by diacylglycerol acyl-CoA transferase (DGAT) activity. The human genome codes for two unrelated enzymes that can perform this function – DGAT1 and DGAT2 (Fig. 4A). DGAT1 is localized in the ER, while DGAT2 is mainly associated with the LDs (48–51). Recently, a DGAT-independent pathway of triglyceride synthesis was described in mammalian cells, but its contribution under normal conditions is believed to be minor, and it was not investigated here (52). Cholesteryl esters are synthesized by acyl-coenzyme A (CoA): cholesterol acyltransferases (ACATs) (Fig. 4A), which in mammalian cells are encoded by related ACAT1 and ACAT2 genes, with ACAT1 representing the major cholesteryl ester synthesizing activity in most cell types (53). To see what pathways of neutral lipid synthesis are upregulated upon the short exposure of cells to PUFAs, we used inhibitors of DGAT1, DGAT2, and ACAT1 and 2, PF-04620110, PF-06424439, and avasimibe, respectively. The experimental design was similar to that in Fig. 3. HeLa cells were incubated in the presence of 200 µM of LCFAs in a normal growth medium (containing serum) for 4 h, and then 0.4 µM of FFA was added for 30 min to monitor the active lipid synthesis pathways. Individual inhibitors were present throughout the whole 4.5 h of incubation in the corresponding samples (Fig. 4B).

**Figure 4.**
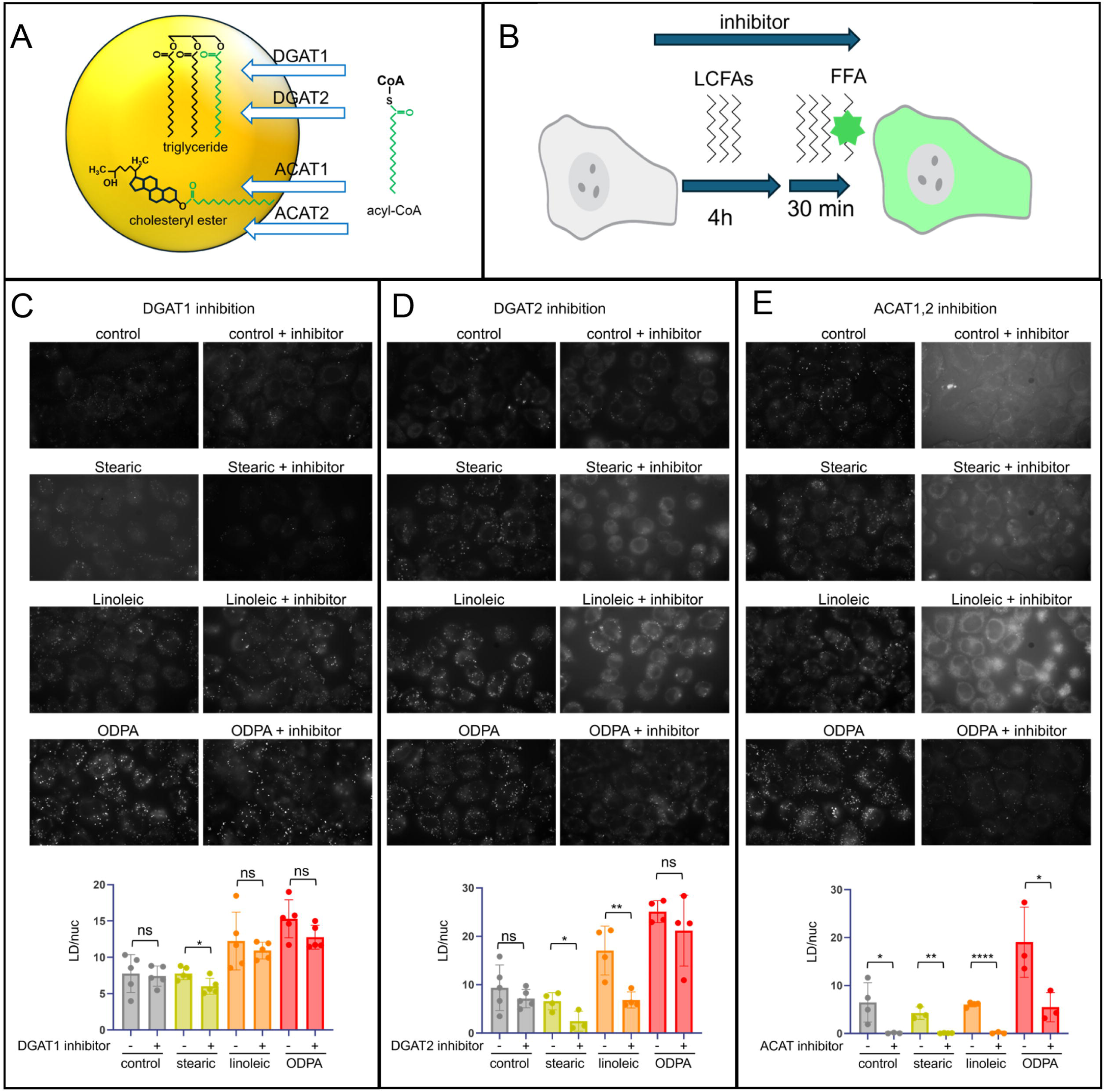
Inhibition of Acyl-CoA cholesterol acyltransferase (ACAT) activity blocks neutral lipid synthesis in PUFA-treated infected cells. **A.** A scheme of LD and the major activities synthesizing triglycerides and cholesterol esters. Green LCFA indicates FFA probe. **B.** A scheme of the experiments used to evaluate the effect of inhibitors on neutral lipid synthesis. **C-D** representative images of live HeLa cells infectedwith poliovirus with an MOI of 50 (or mock-infected controls) and incubated in the presence of 200 µM of the indicated LCFAs for 4 h plus 30 min with the same fatty acids and 0.4 µM of the FFA probe (top) and quantitation of LDs (bottom). Dots represent data for individual randomly chosen fields of view, with at least ∼25 cells per field.

Inhibition of DGAT1 had no significant effect on the number of LDs in either control cells or those incubated with LCFAs, except for a slight but statistically significant decrease of LDs in cells incubated with stearic acid (Fig. 4C). DGAT2 inhibitor was more effective and significantly reduced the number of LDs in cells incubated with stearic (from about 8 to about 1) and linoleic (from ∼15 to ∼6 per cell) acids, with a corresponding increase of the FFA signal in the intracellular membranes, likely reflecting the fatty acid retargeting from inhibited triglyceride to phospholipid synthesis. Yet, its effect on the number of FFA-positive LDs in cells incubated with ODPA was minimal (Fig. 3D). The ACAT inhibitor was the most effective, reducing the number of FFA-positive LDs to an almost undetectable level in control cells and those incubated with stearic or linoleic acids with the corresponding redistribution of the FFA signal to intracellular membranes (Fig. 4E). In cells incubated with ODPA, the number of FFA-positive LDs dropped from ∼20 to ∼5 per cell in the presence of the inhibitor (Fig. 4E). Interestingly, while the number of FFA-positive LDs was significantly reduced, there was no obvious increase of the FFA signal in the intracellular membranes, suggesting that ODPA could strongly outcompete FFA in metabolic reactions supporting the membrane phospholipid synthesis (Fig. 4E).

Thus, inhibition of DGAT2 and especially ACAT activities blocks the targeting of exogenously added LCFAs to LDs and promotes their incorporation into the membranes.

### The anti-viral activity of polyunsaturated fatty acids is increased upon blocking their targeting to lipid droplets and affects multiple steps of the viral replication cycle

To see if preventing PUFAs from being targeted to LDs could increase their anti-viral effect, HeLa cells were infected with poliovirus at an MOI of 50 and incubated for 4 h in normal growth medium with stearic acid (control), linoleic acid, or ODPA, and with or without the ACAT inhibitor. The accumulation of viral protein 2C shows that in the combination of ACAT inhibitor with either linoleic acid or ODPA, the virus-specific signal almost disappeared (Fig. 5A, compare lanes 7 and 2 in both panels). Importantly, while the combination of ACAT inhibitor and ODPA was noticeably toxic for both infected and non-infected cells (as measured by the ATP level), neither infected nor control cells incubated with linoleic acid and the inhibitor showed signs of cytotoxicity (Fig. 5B).

**Figure 5.**
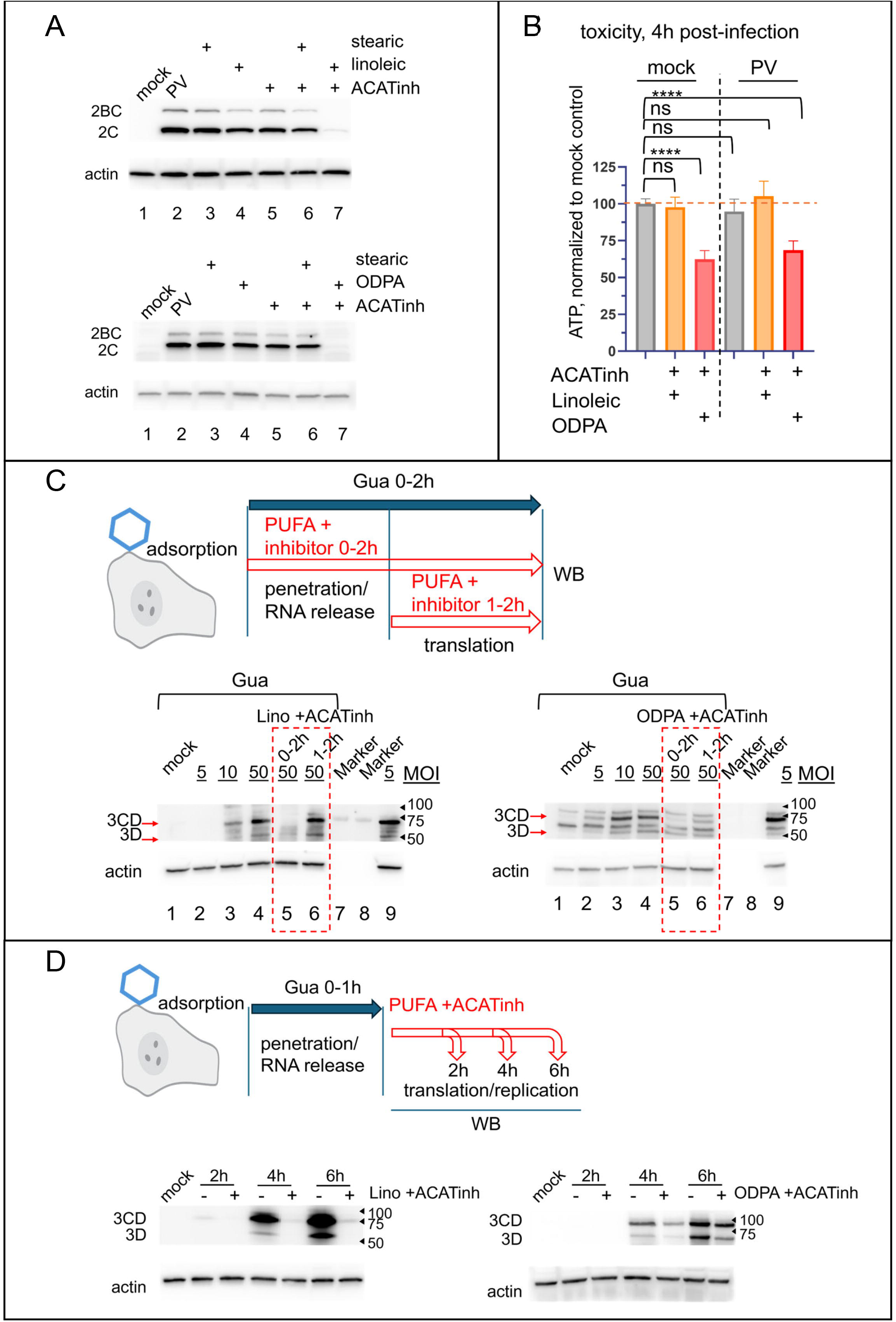
PUFAs can affect multiple steps of the poliovirus life cycle. **A.** Western blots from HeLa cells infected with poliovirus with an MOI of 50 and incubated after virus adsorption with 200 µM of the LCFAs and 50 µM of ACTA inhibitor as indicated for 4 h p.i. **B.** Toxicity assay (ATP level) of cells infected and incubated with LCFAs and ACAT inhibitor as in A. **C.** Top: a scheme of the experiment to evaluate the effect of PUFAs in the presence of ACAT inhibitor on virion penetration/uncoating and initial translation of the viral RNA. Bottom: Western blots of HeLa cells infected with the indicated MOIs (or mock-infected controls) and incubated with (lanes 1-7) or without Gua, an inhibitor of enterovirus replication (lane 9). The indicated PUFA (200 µM) and ACAT inhibitor (50 µM) were added to samples 5 and 6 after virus adsorption or at 1 h p.i., respectively; all cells were lysed at 2 h p.i. **D.** Top: a scheme of the experiment to evaluate the effect of PUFAs in the presence of ACAT inhibitor on the translation/replication step of the viral life cycle. Bottom: Western blots of HeLa cells infected (or mock-infected controls) with an MOI of 50 of poliovirus and incubated for 1 h p.i. in the presence of Gua (an inhibitor of replication) to allow virion penetration and uncoating, and then incubated without Gua in the presence or absence of the indicated PUFA (200 µM) and ACAT inhibitor (50 µM). The cells were lysed at the indicated times post-infection.

Next, we investigated which stage of the viral replication cycle could be affected by PUFA treatment. The replication cycle of a +RNA virus starts with virion attachment, followed by virion internalization and uncoating, leading to the delivery of the genomic RNA to the cytoplasm, where it is translated to produce the first batch of viral proteins required for further development of the replication organelles and establishment of replication complexes. Since the exogenous fatty acids and ACAT inhibitor were added at the beginning of the incubation at 37 °C, after virion adsorption, such treatment could interfere with any stage of the replication cycle, starting from virion internalization.

To see if the compounds affected viral RNA delivery and the initial translation, the cells were incubated in the presence of guanidine-HCl (Gua) for 2 h after virus adsorption, and the level of the viral antigen 3D was assessed by Western blot. Gua is a well-established, strong, and specific inhibitor of enterovirus RNA replication (54, 55), and the potent polyclonal anti-3D antibodies allow the detection of the protein expressed from the input viral RNA, in the absence of replication. The fatty acids with the ACAT inhibitor were added either after virus adsorption together with Gua at the beginning of incubation at 37 °C, or after 1 h of incubation of infected cells with Gua (Fig. 5C). Thus, they were present either from the beginning of the virion penetration step, or after penetration and uncoating is complete, so that the compounds would be able to interfere only with the viral RNA translation happening after 1 h p.i. (Fig. 5C). Additional samples infected with various MOIs were also incubated with Gua as a control for the sensitivity of the system. The 3D signal in samples incubated with Gua for 2 h p.i. was unequivocally detectable starting from an MOI of 10, and was sometimes visible in samples infected with an MOI of 5 (Fig. 5C, lanes 1-4 in both Western blot panels). As expected, Gua effectively blocked viral replication (Fig. 5C, compare lanes 2 and 9 in both panels; these samples were infected with an MOI of 5, but incubated in the presence or absence of Gua, respectively).

The addition of linoleic acid and ACAT inhibitor at the beginning of incubation at 37 °C strongly inhibited the synthesis of viral proteins, while if the compounds were added one hour later, the signal was the same as in the control sample infected with the same MOI (Fig. 5B, left Western blot panel, compare lanes 5, 6, and 4). Thus, the combination of linoleic acid and ACAT inhibitor interferes with the virion penetration and uncoating, but not the translation of viral RNA.

The effect of ODPA and the ACAT inhibitor was similar, although in the latter case, an inhibitory effect during the translation step was also visible (Fig. 5C, right Western blot panel, compare lanes 5, 6, and 4).

To see if the RNA replication step of the viral cycle is affected by PUFA treatment and ACAT inhibition, we infected cells at an MOI of 50 and incubated them at 37 °C for 1 h in the presence of Gua to allow the RNA penetration and initial translation. After 1h, the medium with Gua was removed, and the replication was allowed to proceed in either the presence or absence of the ACAT inhibitor and linoleic acid or ODPA. The cells were lysed at 2, 4, and 6 h p.i., and the amount of the viral antigen 3D was analyzed in a Western blot (Fig. 5D, top panel). ACAT inhibitor and PUFA treatment strongly inhibited viral replication, with the linoleic acid inhibitory effect more pronounced than that of ODPA (Fig. 5D).

Thus, the anti-viral effect of exogenously added PUFAs can include interference with the penetration/uncoating, translation, and replication steps of the enterovirus infectious cycle, and is increased if their targeting to storage in LDs is inhibited.

### Polyunsaturated fatty acids suppress diverse enteroviruses across different cell types with limited potential for resistance

Our data support the model that the anti-viral activity of select PUFAs during the replication step of the enterovirus life cycle is due to their incorporation into the membrane phospholipids, likely making the phospholipid bilayer properties suboptimal for the functioning of the viral replication machinery. Since activation of phospholipid synthesis is a general feature of enterovirus and, at least, some other picornavirus infections (14, 16, 56), these PUFAs should likely have a broad-spectrum anti-viral effect. In addition to poliovirus, which is a member of *Enterovirus coxsackiepol* species, we investigated the effect of linolenic acid and ODPA on Coxsackievirus B3 (CVB3), and Enterovirus D68 (EVD68), representatives of *E. betacoxsakie* and *E. deconjuncti* species, respectively. Infections were performed in HeLa (cervical carcinoma, human, female) and A549 (lung adenocarcinoma, human, male) cells, as representatives of cell types of diverse origin. The cells were infected with an MOI of 10 of these viruses, and incubated in the presence of 200 µM of linoleic acid or ODPA and 50 µM ACAT inhibitor in a normal growth medium (containing serum). Replication of all enteroviruses in both cell cultures was effectively inhibited by the combination of a PUFA and the ACAT inhibitor (Fig 6A).

**Figure 6.**
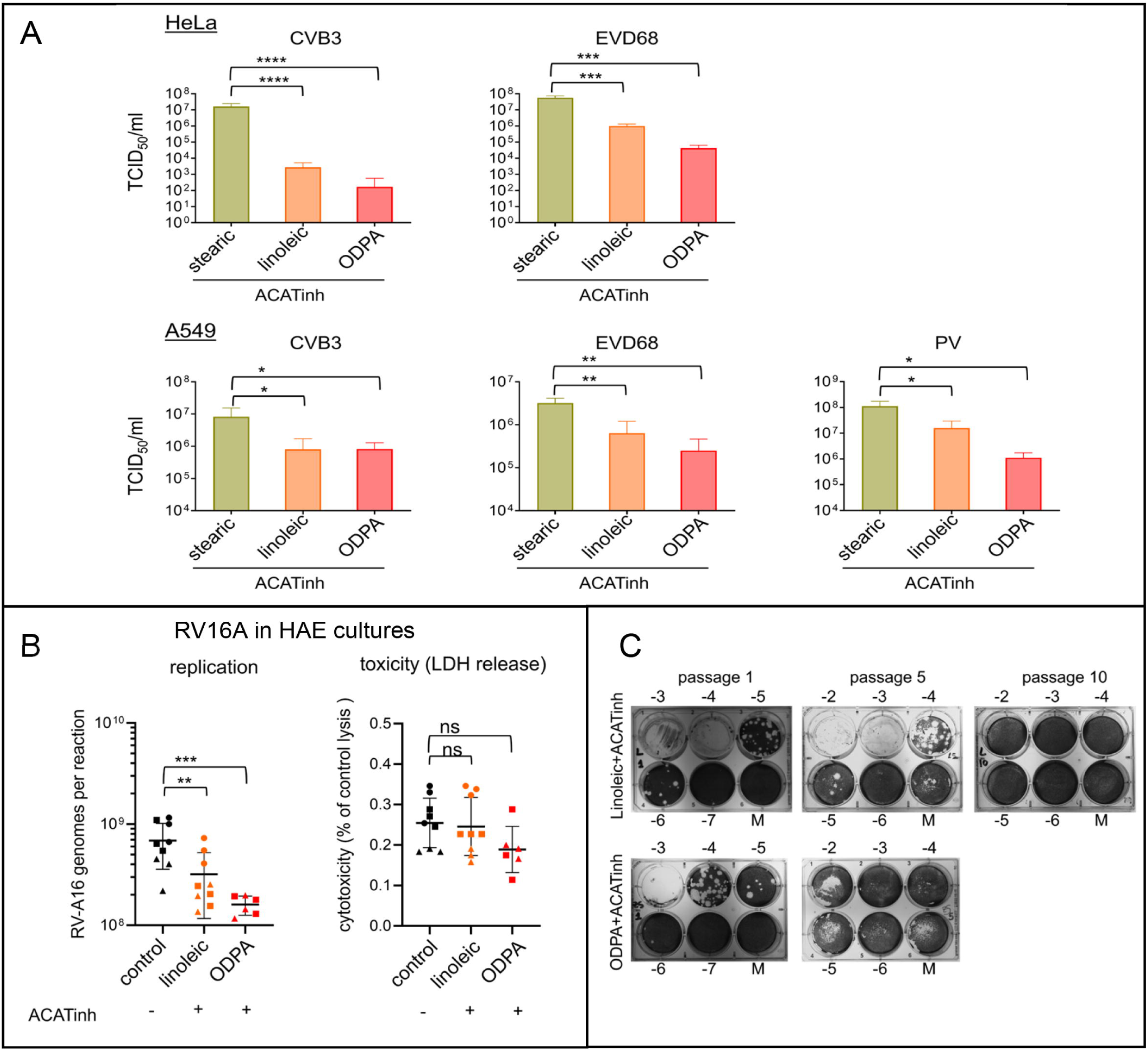
Broad anti-enterovirus activity of select PUFAs is refractory to the emergence of resistance. **A.** HeLa (top) or A549 cells (bottom) were infected with an MOI of 10 of CVB3, poliovirus, or EVD68 and incubated after the virus adsorption in normal growth medium supplemented with 200 µM of stearic acid (control), linoleic acid, or ODPA in the presence of 50 µM of ACAT inhibitor for 6 h (poliovirus) or 8 h (other enteroviruses). Graphs show total virus yield; experiments were performed with three biological replicates. **B.** Differentiated human airway epithelial cultures were infected at the apical surface with RVA16 and incubated for ∼24h in the presence of 50 µM of ACAT inhibitor and 200 µM of either linoleic acid or ODPA (or with a corresponding amount of DMSO vehicle in control). Virus replication was determined by the quantification of intracellular RNA. The basal medium after incubation was used for the toxicity assay (LDH release), in which a lower value indicates less toxicity. Data from cultures differentiated from each donor are indicated by individual symbols. **C.** Poliovirus was serially propagated in HeLa cells in the presence of 200 µM of either linoleic acid or ODPA and 50 µM of ACAT inhibitor. Plaque assays show virus yield from passages one, five, and 10. Note the disappearance of plaques in the ODPA sample from passage five and in the linoleic acid sample from passage 10.

To assess the antiviral potential of LCFAs and ACAT inhibition in a more physiologically relevant system, we tested this treatment on the replication of RVA16 in primary human airway epithelial (HAE) cultures derived from three donors. Following differentiation at the air-liquid interface to mimic respiratory epithelium, HAE cultures were inoculated with RVA16 on the apical surface, and incubated with 50 µM of the ACAT inhibitor along with 200 µM of linoleic acid or ODPA for ∼24 h at 34 °C. To avoid the interference of the original virus inoculum, which in this system can remain trapped in the mucus layer, we washed the apical surface and then assessed the viral replication by measuring the intracellular viral RNA by qPCR. Treatment with LCFAs and ACAT inhibitor was non-toxic in HAE cultures during the prolonged incubation, and virus replication was statistically significantly reduced by the compounds in cultures from all donors (Fig. 6B).

To see if virus resistance can emerge to the LCFA-mediated inhibition of infection, we serially passaged poliovirus in the presence of 200 µM of linoleic acid or ODPA in combination with 50 µM of ACAT inhibitor. For the first passage, the cells were infected at an MOI of 10, and subsequent passages were performed with half of the total virus yield from the previous passage. This scheme usually allows rapid selection of resistance to inhibitors of poliovirus replication (57–59). The material from passages one, five, and ten was analyzed for the presence of infectious virus in a plaque assay. As can be seen from Fig. 6C, virus yield after the first passage was ∼E6 and ∼E5 in the linoleic acid and ODPA samples, respectively. Subsequent passaging in the presence of ODPA and ACAT inhibitor resulted in a complete loss of detectable infectious virus by passage five. The virus passaged in the presence of linoleic acid and ACAT inhibitor was still present at passage five, although the titer decreased compared to the first passage, and the virus was undetectable by passage 10 (Fig. 6C).

Collectively, our data show that LCFAs with specific properties may serve as broad-spectrum inhibitors of enteroviral infections with high barrier to resistance and may represent a promising class of novel anti-viral therapeutics.

## Discussion

The development of effective anti-enterovirus therapeutics so far has proved disappointing. The life cycle of these viruses relies on the generation of a large amount of progeny required for environmental transmission between hosts. The massive genome replication supporting this high virion production depends on an error-prone RNA-dependent RNA polymerase, which inevitably generates a broad spectrum of closely related but distinct variants (quasispecies) that serve as the substrate for effective adaptation to changing conditions, including therapeutic pressure. Resistance is well documented to the prospective anti-enterovirus capsid-targeting compounds pleconaril and V-073 (4), RNA polymerase inhibitor ribavirin (58), 3C protease inhibitors rupintrivir and SG85 (9, 60), as well as to compounds targeting host factors supporting viral replication, such as inhibitors of the activator of small GTPAses Arf GBF1 and a phosphatidylinositol kinase PI4KIIIβ (61, 62). So far, only inhibition of the cellular chaperone Hsp90, involved in the maturation of poliovirus capsid, has been reported to be refractory to the emergence of resistance, but the inhibition of Hsp machinery is toxic to host cells (63).

Here, we explored another strategy, which is not based on inhibition of specific viral or cellular factors supporting the infectious cycle, but rather takes advantage of a major change in the lipid metabolism landscape upon enterovirus infection. The development of membranous replication organelles relies on activation of structural phospholipid synthesis, which in turn consumes LCFAs, which constitute the hydrophobic part of phospholipid molecules. This creates two key shifts in LCFA metabolism in infected *vs.* uninfected cells. First, infected cells require an increased supply of LCFAs. This demand is largely met by activation of the hydrolysis of neutral lipids stored in LDs, but also by the increased import of LCFAs from the extracellular medium due to activation of acyl-CoA synthetases that convert free LCFAs into metabolically active hydrophilic acyl-CoAs. Second, in infected cells, imported LCFAs, which would normally be targeted to neutral lipid synthesis and storage in LDs, are instead incorporated into the membranes of replication organelles (16–18). Thus, feeding infected and non-infected cells with the same LCFA molecules should have different outcomes.

Our results show that, contrary to the original expectations that preferential incorporation of PUFAs into the membranes of replication organelles will promote lipid peroxidation and ferroptosis, the anti-viral activity of LCFAs was largely uncoupled from the degree of unsaturation and the propensity to induce ferroptosis. Rather, PUFA molecules of specific shapes were incompatible with viral replication. While we explored only a small selection of natural LCFAs, our observations open the possibility of designing new LCFA-like molecules with improved stability in an anti-viral conformation. Of course, the balance between the anti-viral effect and the toxicity to non-infected cells should be an important consideration for the development of LCFA-based anti-virals. In our proof-of-concept studies, we used an inhibitor of cholesteryl ester synthesis to promote LCFA incorporation into phospholipids. A combination of this inhibitor with a highly polyunsaturated LCFA could be toxic for both control and infected cells during prolonged incubation, while in the absence of the inhibitor, cells could easily manage incubation with a high concentration of PUFAs, suggesting that an acceptable selectivity index of treatment with both compounds could be achieved. Optimized LCFA-based anti-viral molecules should retain the capacity of being recognized by acyl-CoA synthetases and acyl-CoA transferases, preferentially those that are active in infected but not in uninfected cells. Unfortunately, current knowledge of the profiles of expression and activity of these enzymes in cells relevant for enterovirus replication is very limited.

An important advantage of LCFA-mediated anti-viral activity is that it is refractory to the emergence of resistant mutants. We applied a selection scheme that usually results in a rapid selection of poliovirus mutants resistant to replication inhibitors, yet no mutants capable of replicating under conditions of increased supply of specific LCFAs emerged. This likely reflects their multi-pronged effect on the infection cycle, associated with the incorporation of these LCFAs into membranes. The altered membrane properties likely interfere with the activity of multiple viral as well as cellular factors required to support the infection, which is impossible to overcome with a few point mutations. More extensive studies of the development of resistance in physiologically relevant conditions will be required in the future. If resistant mutants can be selected, they may reveal what viral proteins are specifically affected by the changes in membrane phospholipid composition.

In the absence of an inhibitor of neutral lipid synthesis, the anti-viral effect of PUFAs in cell culture in the presence of serum on one replication cycle was moderate, within half to one log_10_ of virus yield. The inhibition of neutral lipid synthesis significantly enhanced the inhibitory effect, consistent with increased incorporation of the LCFAs into membranes of replication organelles. Interestingly, in a more physiologically relevant model of human airway epithelium system, treatment with LCFAs in combination with the ACAT inhibitor was much better tolerated than in cell lines during prolonged incubation, but the anti-viral effect was not as dramatic. This could reflect a different profile of expression of enzymes involved in neutral and phospholipid synthesis in differentiated cells compared to the continuous cell lines.

Even a moderate sustained reduction of viral replication may be sufficient for the prevention of clinical symptoms and disease progression. It is worth noting that the propagation yield of the poliovirus attenuated vaccine Sabin I strain in normal conditions in HeLa cells was reported to be either comparable or less than half a log_10_ of that of the virulent strain Mahoney (64, 65). Our results show that LCFAs that are part of a normal diet, such as linoleic acid, can suppress replication of diverse enteroviuses, suggesting that the composition of dietary lipids may determine the outcome of enterovirus infections. Whether pathological complications of enterovirus infections, which are normally mild and self-resolving, can be linked to a specific nutritional state of the host could be an interesting research direction.

Overall, our data show that infection-specific metabolic rewiring can be harnessed to develop anti-viral strategies that do not rely on inhibiting specific viral or host proteins. In particular, they highlight the potential of LCFAs and their metabolism for the development of broadly effective inhibitors of enterovirus replication with a high barrier to the development of resistance.

## Acknowledgements

The work was supported by the NIH R01 AI169458 grant to GAB and MAS. AZ was supported by the Basil & Anne Hatziolos Scholarship Fund.

---

**Table S1.** RDKit-derived conformational and shape parameters for long-chain fatty acids.

Reported values summarize the conformer ensemble for each compound. n_conformers indicates the number of valid conformers included in the analysis. L_mean, W_mean, and T_mean are the means of the molecular length, width, and thickness across conformers, respectively. EndToEnd_mean is the mean distance between the terminal atoms defining the long-chain axis. Rg_mean is the mean radius of gyration, describing overall spatial compactness. NPR1_mean and NPR2_mean are the mean normalized principal moments of inertia, which describe molecular shape anisotropy, with values reflecting rod-like, disk-like, or more spherical conformations. V_vdw_mean is the mean van der Waals molecular volume. V_sas_p1p4_mean is the mean solvent-accessible surface-derived volume calculated using a 1.4 Å probe radius. The columns with _sd report the standard deviation of the corresponding values

